## Supplemental materials and figures for "Mouse vendor influence on the bacterial and viral gut composition exceeds the effect of diet": Rasmussen et al. 2019 - Viruses - SM.docx

**Table S1.** Illumina NextSeq sequencing details of both the 16S rRNA gene amplicons and metaviromes of faecal content isolated from both cecum and colon. In addition, listing the amount of b- and vOTUs after filtering as described in methods. BC = bacterial community, VC = viral community.

|  | **BC** | | **VC** | |
| --- | --- | --- | --- | --- |
|  | **Cecum** | **Colon** | **Cecum** | **Colon** |
| Mean sequencing depth (reads) | 318395 | 168388 | 829533 | 456452 |
| STD (reads) | 173421 | 28037 | 336144 | 110063 |
| Lowest sequencing depth (reads) | 47182 | 47004 | 212545 | 63183 |
| Maximum sequencing depth (reads) | 808971 | 223787 | 1621360 | 643913 |
| bOTU/vOTUs before filtering | 6205 | 6205 | 12624 | 12624 |
| bOTU/vOTUs after filtering | 2653 | 3259 | 3441 | 2073 |

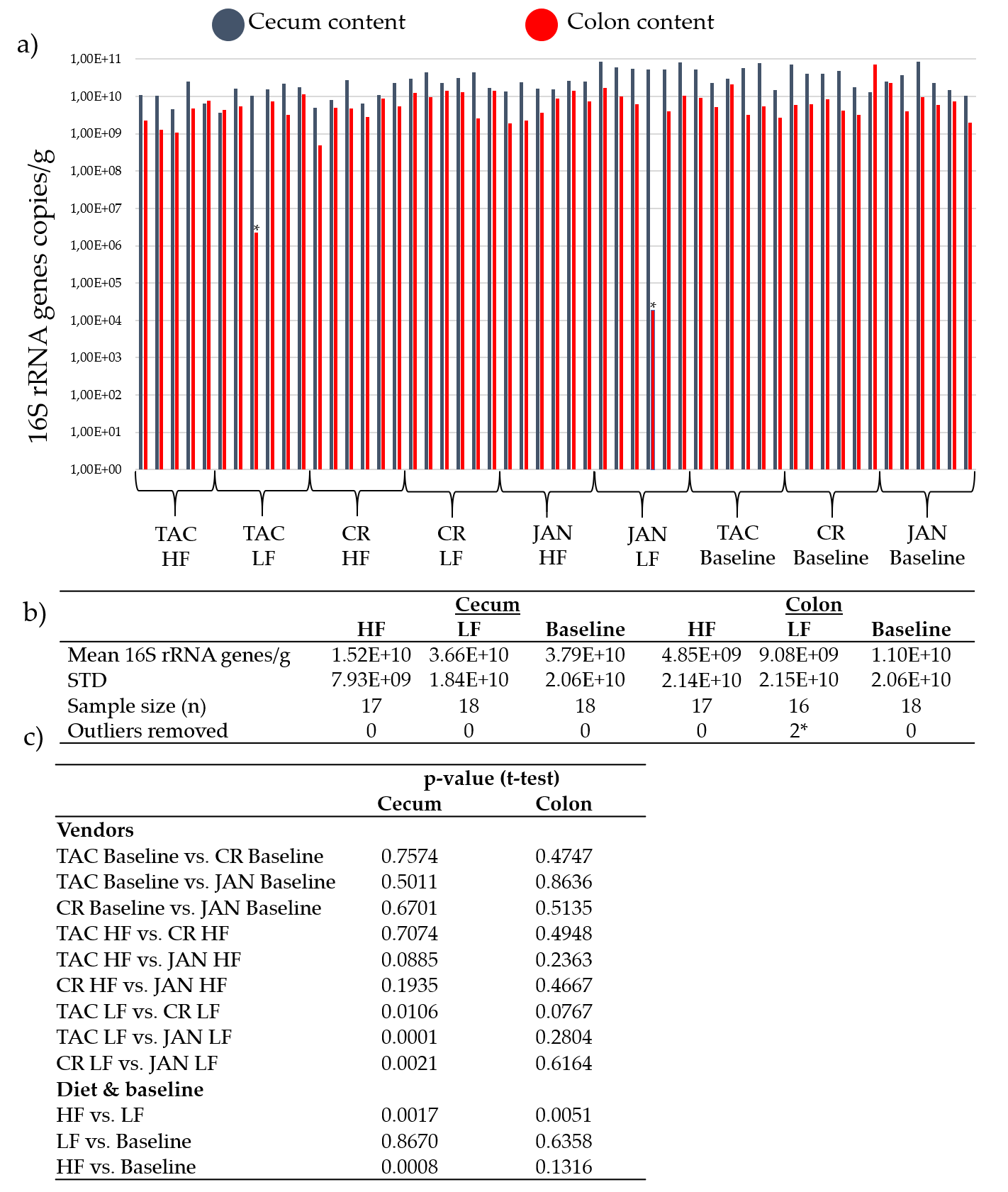

**Figure S1.** a) Mean qPCR results of universal primers targeting 16S rRNA genes (V3 region as described in methods) to estimate bacterial density in the included samples. All samples are included in the bar plot to show sample variations. b) Mean 16S rRNA gene copies/g in cecum and colon samples in the HF, LF and baseline groups. c) A t-test of differences in mean 16S rRNA gene copies/g between HF, LF, baseline, and between vendors. HF diet = high-fat, LF = low-fat diet, CR = Charles River, JAN = Janvier, TAC = Taconic. * sample 34 and 134 were removed as outliers.

### Figure S2: Bacterial and viral α-diversity analysis with other indices – Cecum

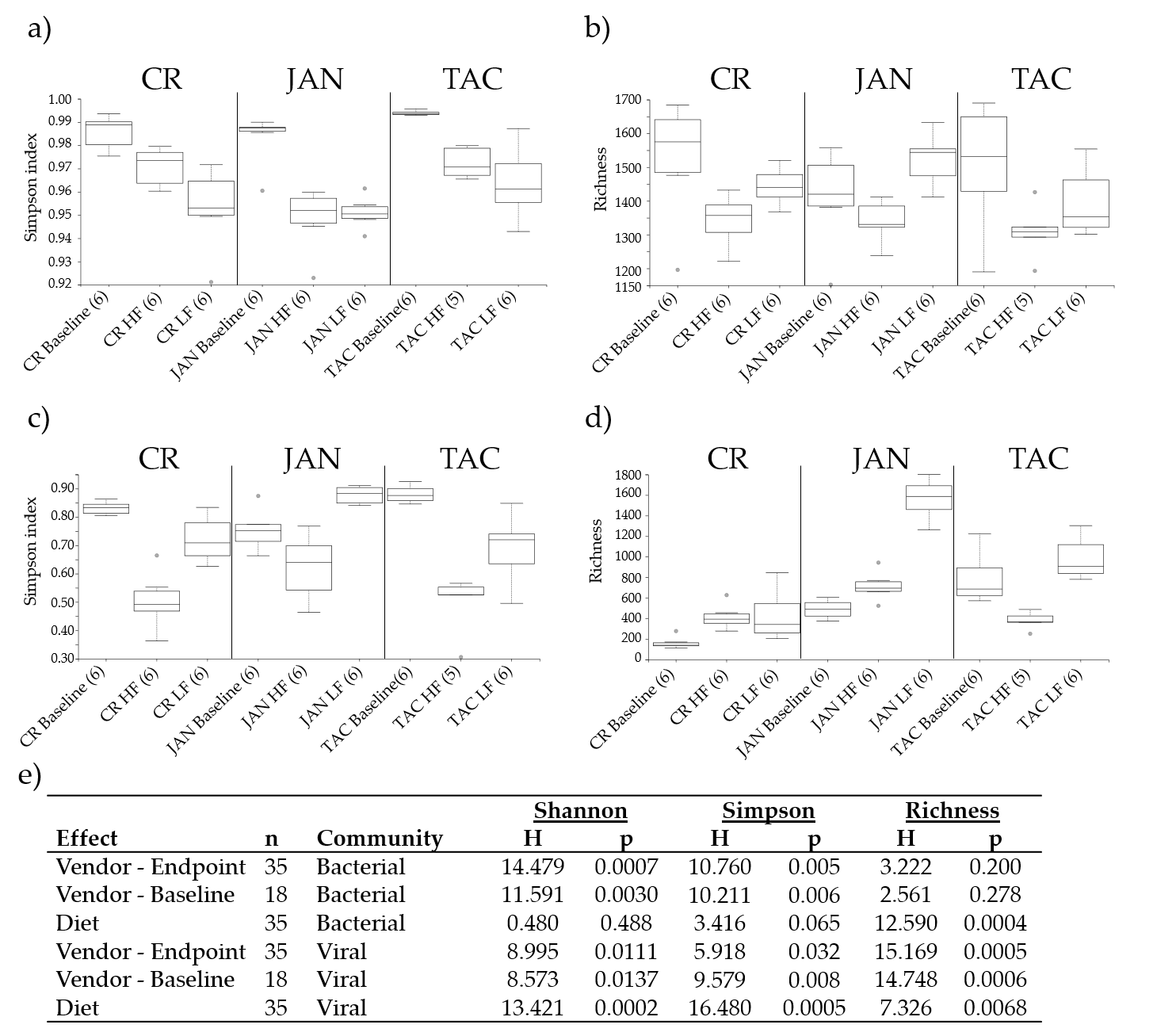

**Figure S2.** α-diversity (Simpson index and Richness) of the caecal bacterial community in a) and b), and viral community in c) and d). The parentheses show the number of samples from each group included in the plot. e) Kruskal Wallis group analysis of the α-diversity indices of the effects of diet and vendor at baseline and endpoint (18 weeks of age). Abbreviations: LF = low-fat diet, HF = high-fat diet, CR = Charles River, JAN = Janvier, TAC = Taconic.

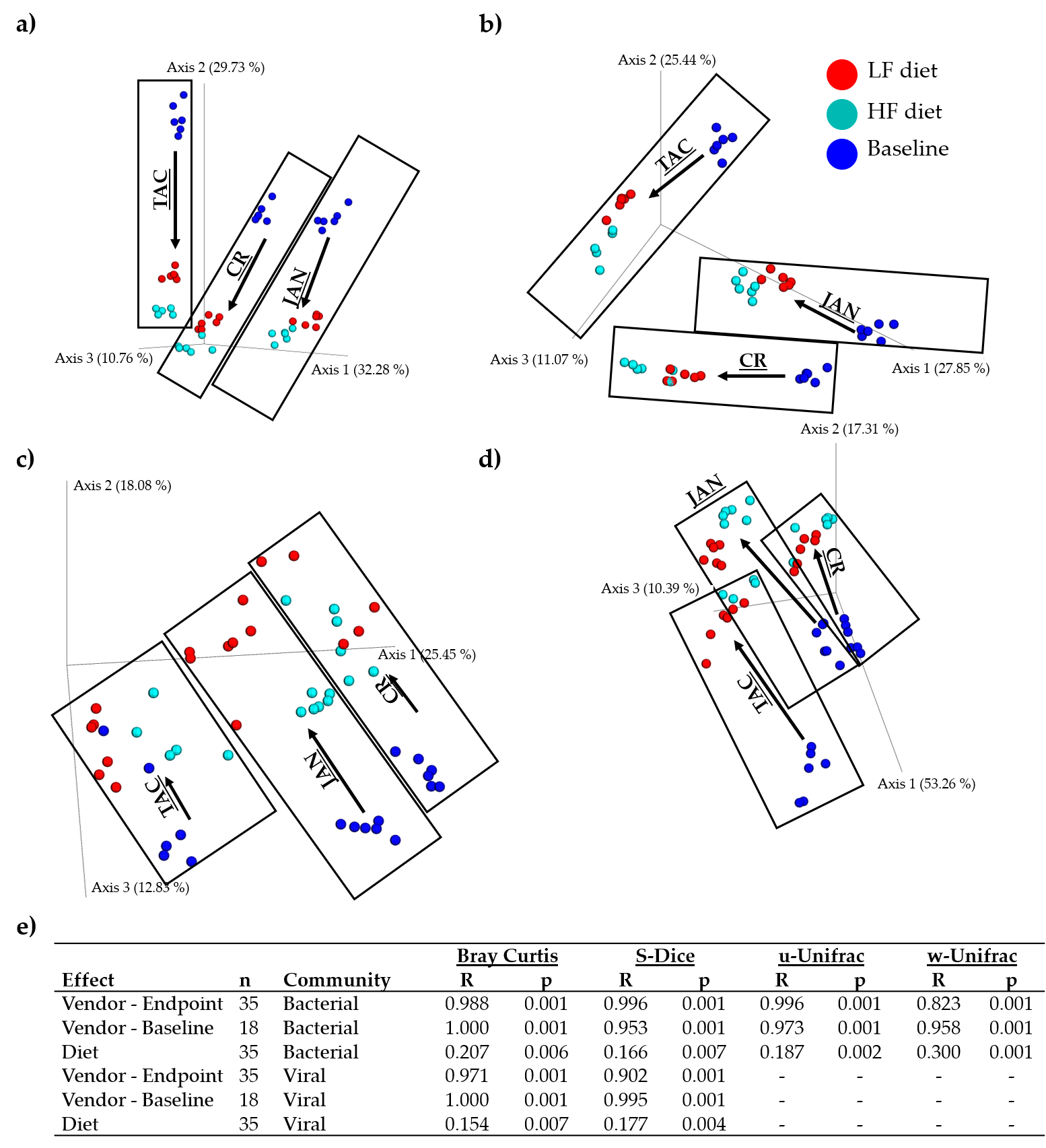

**Figure S3.** β-diversity (Sørensen-Dice, unweighted- and weighted unifrac metrics) of the caecal bacterial community in a), b), and d), and viral community in c) illustrated with PCoA plots. e) ANOSIM of the effects of diet and vendor at baseline and endpoint (18 weeks of age). CR = Charles River, JAN = Janvier, TAC = Taconic. Black boxes frame the samples associated to the mice vendor. S-Dice = Sørensen-Dice, u = unweighted, w = weighted.

**Table S1.** Pairwise Kruskal Wallis (H and p-values) comparison of α-diversity of both the caecal bacterial (BC) and viral community (VC) of all applied indices. BL = baseline, HF = high-fat diet, LF = low-fat diet.

|  |  |  | **Shannon-BC** | | **Shannon-VC** | | **Simpson-BC** | | **Simpson -VC** | | **Richness-BC** | | **Richness-VC** | |
| --- | --- | --- | --- | --- | --- | --- | --- | --- | --- | --- | --- | --- | --- | --- |
| **Gr. 1** | **Gr. 2** | **n** | **H** | **p** | **H** | **p** | **H** | **p** | **H** | **p** | **H** | **p** | **H** | **p** |
| CR_BL | CR_HF | 12 | 6.564 | 0.010 | 6.564 | 0.010 | 5.026 | 0.025 | 8.308 | 0.004 | 3.692 | 0.055 | 7.880 | 0.005 |
|  | CR_LF | 12 | 8.308 | 0.004 | 1.256 | 0.262 | 8.308 | 0.004 | 5.026 | 0.025 | 2.077 | 0.150 | 6.564 | 0.010 |
|  | JAN_BL | 12 | 1.641 | 0.200 | 0.000 | 1.000 | 0.923 | 0.337 | 3.692 | 0.055 | 2.077 | 0.150 | 8.308 | 0.004 |
|  | JAN_HF | 12 | 8.308 | 0.004 | 0.923 | 0.337 | 8.308 | 0.004 | 8.308 | 0.004 | 3.692 | 0.055 | 8.308 | 0.004 |
|  | JAN_LF | 12 | 8.308 | 0.004 | 8.308 | 0.004 | 8.308 | 0.004 | 5.026 | 0.025 | 0.410 | 0.522 | 8.308 | 0.004 |
|  | TAC_BL | 12 | 7.410 | 0.006 | 8.308 | 0.004 | 5.769 | 0.016 | 5.769 | 0.016 | 0.026 | 0.873 | 8.308 | 0.004 |
|  | TAC_HF | 11 | 7.500 | 0.006 | 7.500 | 0.006 | 4.033 | 0.045 | 7.500 | 0.006 | 4.033 | 0.045 | 6.533 | 0.011 |
|  | TAC_LF | 12 | 5.769 | 0.016 | 0.103 | 0.749 | 6.564 | 0.010 | 5.026 | 0.025 | 2.077 | 0.150 | 8.308 | 0.004 |
| CR_HF | CR_LF | 12 | 4.333 | 0.037 | 4.333 | 0.037 | 5.026 | 0.025 | 6.564 | 0.010 | 5.769 | 0.016 | 0.231 | 0.631 |
|  | JAN_BL | 12 | 5.026 | 0.025 | 6.564 | 0.010 | 4.333 | 0.037 | 7.410 | 0.006 | 1.641 | 0.200 | 1.256 | 0.262 |
|  | JAN_HF | 12 | 8.308 | 0.004 | 3.692 | 0.055 | 8.308 | 0.004 | 3.103 | 0.078 | 0.026 | 0.873 | 7.410 | 0.006 |
|  | JAN_LF | 12 | 6.564 | 0.010 | 8.308 | 0.004 | 6.564 | 0.010 | 8.308 | 0.004 | 7.410 | 0.006 | 8.308 | 0.004 |
|  | TAC_BL | 12 | 8.308 | 0.004 | 8.308 | 0.004 | 8.308 | 0.004 | 8.308 | 0.004 | 3.103 | 0.078 | 6.182 | 0.013 |
|  | TAC_HF | 11 | 0.533 | 0.465 | 0.033 | 0.855 | 0.133 | 0.715 | 0.300 | 0.584 | 0.533 | 0.465 | 0.133 | 0.715 |
|  | TAC_LF | 12 | 0.026 | 0.873 | 5.026 | 0.025 | 0.923 | 0.337 | 5.026 | 0.025 | 0.410 | 0.522 | 8.308 | 0.004 |
| CR_LF | JAN_BL | 12 | 7.410 | 0.006 | 0.641 | 0.423 | 6.564 | 0.010 | 0.641 | 0.423 | 0.006 | 0.936 | 0.923 | 0.337 |
|  | JAN_HF | 12 | 2.564 | 0.109 | 0.000 | 1.000 | 0.231 | 0.631 | 2.077 | 0.150 | 5.769 | 0.016 | 3.692 | 0.055 |
|  | JAN_LF | 12 | 0.923 | 0.337 | 8.308 | 0.004 | 0.641 | 0.423 | 8.308 | 0.004 | 3.692 | 0.055 | 8.308 | 0.004 |
|  | TAC_BL | 12 | 8.308 | 0.004 | 4.333 | 0.037 | 8.308 | 0.004 | 8.308 | 0.004 | 0.923 | 0.337 | 4.333 | 0.037 |
|  | TAC_HF | 11 | 2.700 | 0.100 | 7.500 | 0.006 | 3.333 | 0.068 | 7.500 | 0.006 | 5.633 | 0.018 | 0.033 | 0.855 |
|  | TAC_LF | 12 | 2.564 | 0.109 | 0.103 | 0.749 | 0.923 | 0.337 | 0.026 | 0.873 | 1.256 | 0.262 | 6.564 | 0.010 |
| JAN_BL | JAN_HF | 12 | 8.308 | 0.004 | 0.641 | 0.423 | 8.308 | 0.004 | 4.333 | 0.037 | 1.641 | 0.200 | 5.769 | 0.016 |
|  | JAN_LF | 12 | 8.308 | 0.004 | 7.410 | 0.006 | 7.410 | 0.006 | 5.769 | 0.016 | 2.564 | 0.109 | 8.308 | 0.004 |
|  | TAC_BL | 12 | 8.308 | 0.004 | 4.333 | 0.037 | 8.308 | 0.004 | 5.769 | 0.016 | 1.641 | 0.200 | 7.410 | 0.006 |
|  | TAC_HF | 11 | 5.633 | 0.018 | 7.500 | 0.006 | 3.333 | 0.068 | 7.500 | 0.006 | 2.133 | 0.144 | 3.333 | 0.068 |
|  | TAC_LF | 12 | 5.769 | 0.016 | 0.103 | 0.749 | 5.026 | 0.025 | 0.923 | 0.337 | 0.641 | 0.423 | 8.308 | 0.004 |
| JAN_HF | JAN_LF | 12 | 0.410 | 0.522 | 8.308 | 0.004 | 0.000 | 1.000 | 8.308 | 0.004 | 7.880 | 0.005 | 8.308 | 0.004 |
|  | TAC_BL | 12 | 8.308 | 0.004 | 4.333 | 0.037 | 8.308 | 0.004 | 8.308 | 0.004 | 3.103 | 0.078 | 0.000 | 1.000 |
|  | TAC_HF | 11 | 7.500 | 0.006 | 2.700 | 0.100 | 7.500 | 0.006 | 2.700 | 0.100 | 0.833 | 0.361 | 7.500 | 0.006 |
|  | TAC_LF | 12 | 5.769 | 0.016 | 0.231 | 0.631 | 2.077 | 0.150 | 1.256 | 0.262 | 0.410 | 0.522 | 5.026 | 0.025 |
| JAN_LF | TAC_BL | 12 | 8.308 | 0.004 | 6.564 | 0.010 | 8.308 | 0.004 | 0.026 | 0.873 | 0.026 | 0.873 | 8.308 | 0.004 |
|  | TAC_HF | 11 | 5.633 | 0.018 | 7.500 | 0.006 | 7.500 | 0.006 | 7.500 | 0.006 | 6.533 | 0.011 | 7.500 | 0.006 |
|  | TAC_LF | 12 | 5.026 | 0.025 | 6.564 | 0.010 | 3.103 | 0.078 | 6.564 | 0.010 | 3.692 | 0.055 | 7.410 | 0.006 |
| TAC_BL | TAC_HF | 11 | 7.500 | 0.006 | 7.500 | 0.006 | 7.500 | 0.006 | 7.500 | 0.006 | 2.700 | 0.100 | 7.500 | 0.006 |
|  | TAC_LF | 12 | 8.308 | 0.004 | 1.641 | 0.200 | 8.308 | 0.004 | 7.410 | 0.006 | 1.641 | 0.200 | 2.077 | 0.150 |
| TAC_HF | TAC_LF | 11 | 0.133 | 0.715 | 7.500 | 0.006 | 1.633 | 0.201 | 4.033 | 0.045 | 2.133 | 0.144 | 7.500 | 0.006 |

**Table S3.** Pairwise ANOSIM (R and p-values) comparison of β-diversity of both the caecal bacterial (BC) and viral community (VC) of all applied metrices. The R-value is an arbitrary level of separation. S-Dice = Sørensen-Dice, u = unweighted, w = weighted.

|  |  |  | **Bray-Curtis-BC** | | **Bray-Curtis-VC** | | **S-Dice-BC** | | **S-Dice-VC** | | **u-UniFrac-BC** | | **w-UniFrac-BC** | |
| --- | --- | --- | --- | --- | --- | --- | --- | --- | --- | --- | --- | --- | --- | --- |
| **Gr. 1** | **Gr. 2** | **n** | **R** | **p** | **R** | **p** | **R** | **p** | **R** | **p** | **R** | **p** | **R** | **p** |
| CR_BL | CR_HF | 12 | 1.000 | 0.003 | 1.000 | 0.003 | 1.000 | 0.005 | 0.937 | 0.001 | 1.000 | 0.003 | 1.000 | 0.001 |
|  | CR_LF | 12 | 1.000 | 0.005 | 1.000 | 0.001 | 1.000 | 0.003 | 0.970 | 0.002 | 1.000 | 0.004 | 1.000 | 0.004 |
|  | JAN_BL | 12 | 1.000 | 0.002 | 1.000 | 0.006 | 1.000 | 0.002 | 0.987 | 0.005 | 0.990 | 0.003 | 0.830 | 0.004 |
|  | JAN_HF | 12 | 1.000 | 0.001 | 1.000 | 0.005 | 1.000 | 0.006 | 0.996 | 0.005 | 1.000 | 0.002 | 1.000 | 0.003 |
|  | JAN_LF | 12 | 1.000 | 0.003 | 1.000 | 0.003 | 1.000 | 0.001 | 1.000 | 0.003 | 1.000 | 0.002 | 1.000 | 0.003 |
|  | TAC_BL | 12 | 1.000 | 0.001 | 1.000 | 0.002 | 1.000 | 0.002 | 1.000 | 0.001 | 1.000 | 0.004 | 1.000 | 0.003 |
|  | TAC_HF | 11 | 1.000 | 0.003 | 1.000 | 0.003 | 1.000 | 0.004 | 1.000 | 0.004 | 1.000 | 0.005 | 1.000 | 0.003 |
|  | TAC_LF | 12 | 1.000 | 0.002 | 1.000 | 0.003 | 1.000 | 0.002 | 1.000 | 0.006 | 1.000 | 0.002 | 1.000 | 0.006 |
| CR_HF | CR_LF | 12 | 0.980 | 0.002 | 0.652 | 0.004 | 0.850 | 0.003 | 0.344 | 0.031 | 0.850 | 0.001 | 0.650 | 0.003 |
|  | JAN_BL | 12 | 1.000 | 0.001 | 1.000 | 0.006 | 1.000 | 0.004 | 1.000 | 0.006 | 1.000 | 0.002 | 1.000 | 0.002 |
|  | JAN_HF | 12 | 1.000 | 0.001 | 1.000 | 0.002 | 1.000 | 0.003 | 0.985 | 0.002 | 1.000 | 0.004 | 0.970 | 0.003 |
|  | JAN_LF | 12 | 1.000 | 0.002 | 1.000 | 0.001 | 1.000 | 0.002 | 0.989 | 0.004 | 1.000 | 0.005 | 1.000 | 0.002 |
|  | TAC_BL | 12 | 1.000 | 0.003 | 1.000 | 0.003 | 1.000 | 0.002 | 1.000 | 0.005 | 1.000 | 0.003 | 1.000 | 0.002 |
|  | TAC_HF | 11 | 0.980 | 0.001 | 1.000 | 0.004 | 1.000 | 0.004 | 1.000 | 0.001 | 1.000 | 0.004 | 0.800 | 0.001 |
|  | TAC_LF | 12 | 1.000 | 0.002 | 1.000 | 0.004 | 1.000 | 0.003 | 1.000 | 0.003 | 1.000 | 0.003 | 1.000 | 0.003 |
| CR_LF | JAN_BL | 12 | 1.000 | 0.004 | 1.000 | 0.003 | 1.000 | 0.002 | 0.998 | 0.006 | 1.000 | 0.003 | 1.000 | 0.004 |
|  | JAN_HF | 12 | 1.000 | 0.004 | 1.000 | 0.003 | 1.000 | 0.001 | 0.928 | 0.004 | 1.000 | 0.005 | 1.000 | 0.002 |
|  | JAN_LF | 12 | 1.000 | 0.003 | 0.961 | 0.002 | 1.000 | 0.005 | 0.802 | 0.002 | 1.000 | 0.003 | 1.000 | 0.003 |
|  | TAC_BL | 12 | 1.000 | 0.005 | 1.000 | 0.003 | 1.000 | 0.001 | 0.970 | 0.001 | 1.000 | 0.002 | 1.000 | 0.005 |
|  | TAC_HF | 11 | 1.000 | 0.002 | 1.000 | 0.004 | 1.000 | 0.005 | 0.984 | 0.002 | 1.000 | 0.007 | 0.970 | 0.005 |
|  | TAC_LF | 12 | 1.000 | 0.003 | 1.000 | 0.003 | 1.000 | 0.003 | 0.961 | 0.002 | 1.000 | 0.002 | 1.000 | 0.002 |
| JAN_BL | JAN_HF | 12 | 1.000 | 0.004 | 1.000 | 0.003 | 1.000 | 0.004 | 1.000 | 0.003 | 1.000 | 0.003 | 1.000 | 0.002 |
|  | JAN_LF | 12 | 1.000 | 0.002 | 1.000 | 0.004 | 1.000 | 0.006 | 1.000 | 0.001 | 1.000 | 0.004 | 1.000 | 0.002 |
|  | TAC_BL | 12 | 1.000 | 0.003 | 1.000 | 0.003 | 1.000 | 0.001 | 1.000 | 0.002 | 1.000 | 0.003 | 1.000 | 0.005 |
|  | TAC_HF | 11 | 1.000 | 0.003 | 1.000 | 0.002 | 1.000 | 0.003 | 1.000 | 0.005 | 1.000 | 0.001 | 1.000 | 0.002 |
|  | TAC_LF | 12 | 1.000 | 0.004 | 1.000 | 0.003 | 1.000 | 0.003 | 1.000 | 0.002 | 1.000 | 0.004 | 1.000 | 0.003 |
| JAN_HF | JAN_LF | 12 | 0.980 | 0.003 | 0.967 | 0.004 | 0.990 | 0.002 | 0.961 | 0.001 | 0.990 | 0.003 | 1.000 | 0.004 |
|  | TAC_BL | 12 | 1.000 | 0.002 | 1.000 | 0.002 | 1.000 | 0.001 | 1.000 | 0.002 | 1.000 | 0.005 | 1.000 | 0.002 |
|  | TAC_HF | 11 | 1.000 | 0.001 | 1.000 | 0.005 | 1.000 | 0.002 | 1.000 | 0.002 | 1.000 | 0.002 | 0.980 | 0.002 |
|  | TAC_LF | 12 | 1.000 | 0.003 | 1.000 | 0.004 | 1.000 | 0.005 | 1.000 | 0.004 | 1.000 | 0.003 | 1.000 | 0.004 |
| JAN_LF | TAC_BL | 12 | 1.000 | 0.005 | 1.000 | 0.002 | 1.000 | 0.004 | 0.996 | 0.002 | 1.000 | 0.003 | 1.000 | 0.004 |
|  | TAC_HF | 11 | 1.000 | 0.001 | 1.000 | 0.003 | 1.000 | 0.001 | 1.000 | 0.004 | 1.000 | 0.004 | 1.000 | 0.002 |
|  | TAC_LF | 12 | 1.000 | 0.002 | 1.000 | 0.002 | 1.000 | 0.002 | 1.000 | 0.004 | 1.000 | 0.005 | 1.000 | 0.001 |
| TAC_BL | TAC_HF | 11 | 1.000 | 0.004 | 1.000 | 0.001 | 1.000 | 0.002 | 0.973 | 0.003 | 1.000 | 0.006 | 1.000 | 0.002 |
|  | TAC_LF | 12 | 1.000 | 0.004 | 1.000 | 0.006 | 1.000 | 0.002 | 0.959 | 0.003 | 1.000 | 0.003 | 1.000 | 0.006 |
| TAC_HF | TAC_LF | 11 | 0.880 | 0.006 | 0.963 | 0.005 | 0.970 | 0.003 | 0.824 | 0.005 | 0.990 | 0.003 | 0.570 | 0.001 |

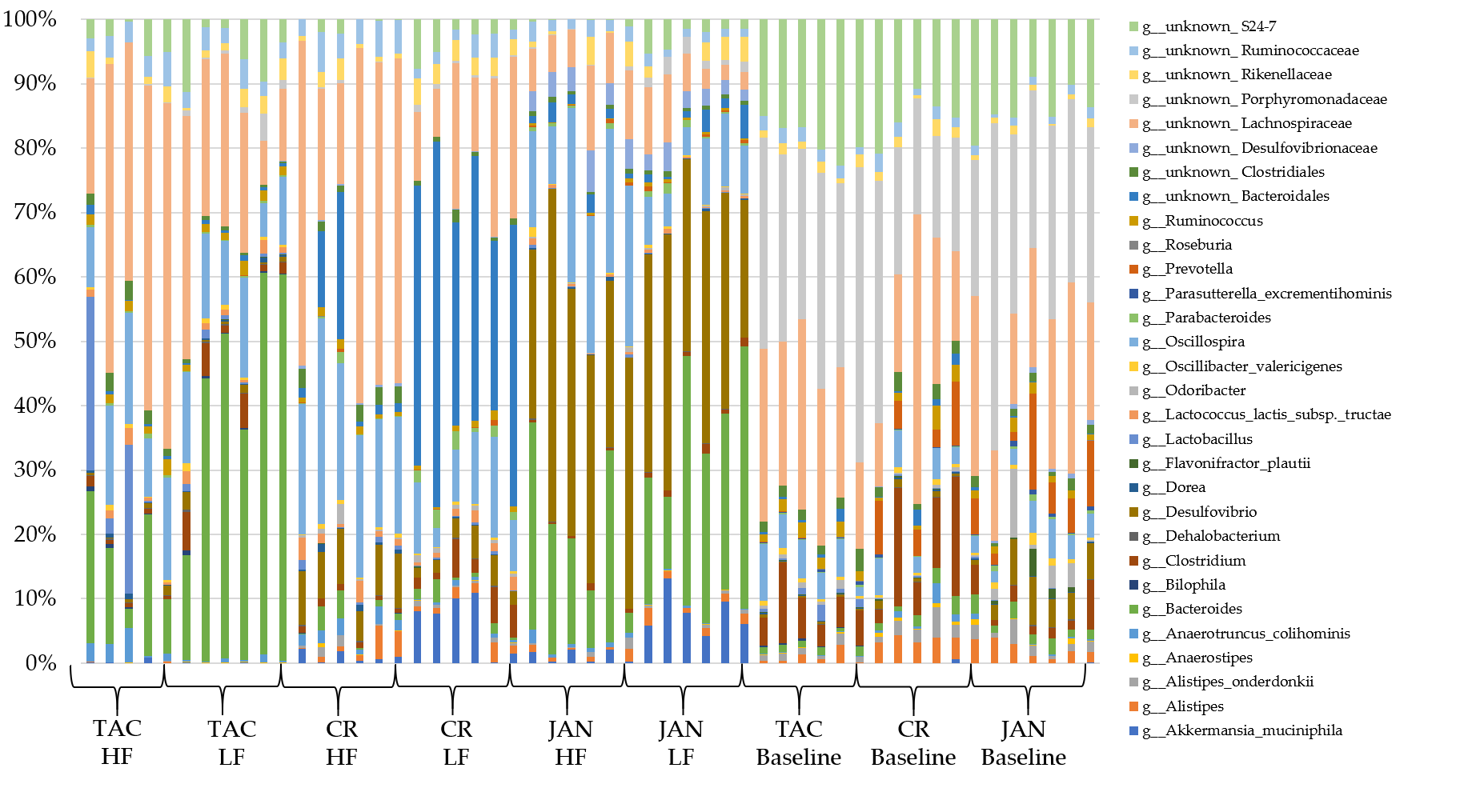

**Figure S4.** Bacterial taxonomy on genus level of all individual cecum samples. A threshold of the relative abundance was set to 0.25%. HF = high-fat diet, LF = low-fat diet. TAC = Taconic, CR = Charles River, JAN = Janvier.

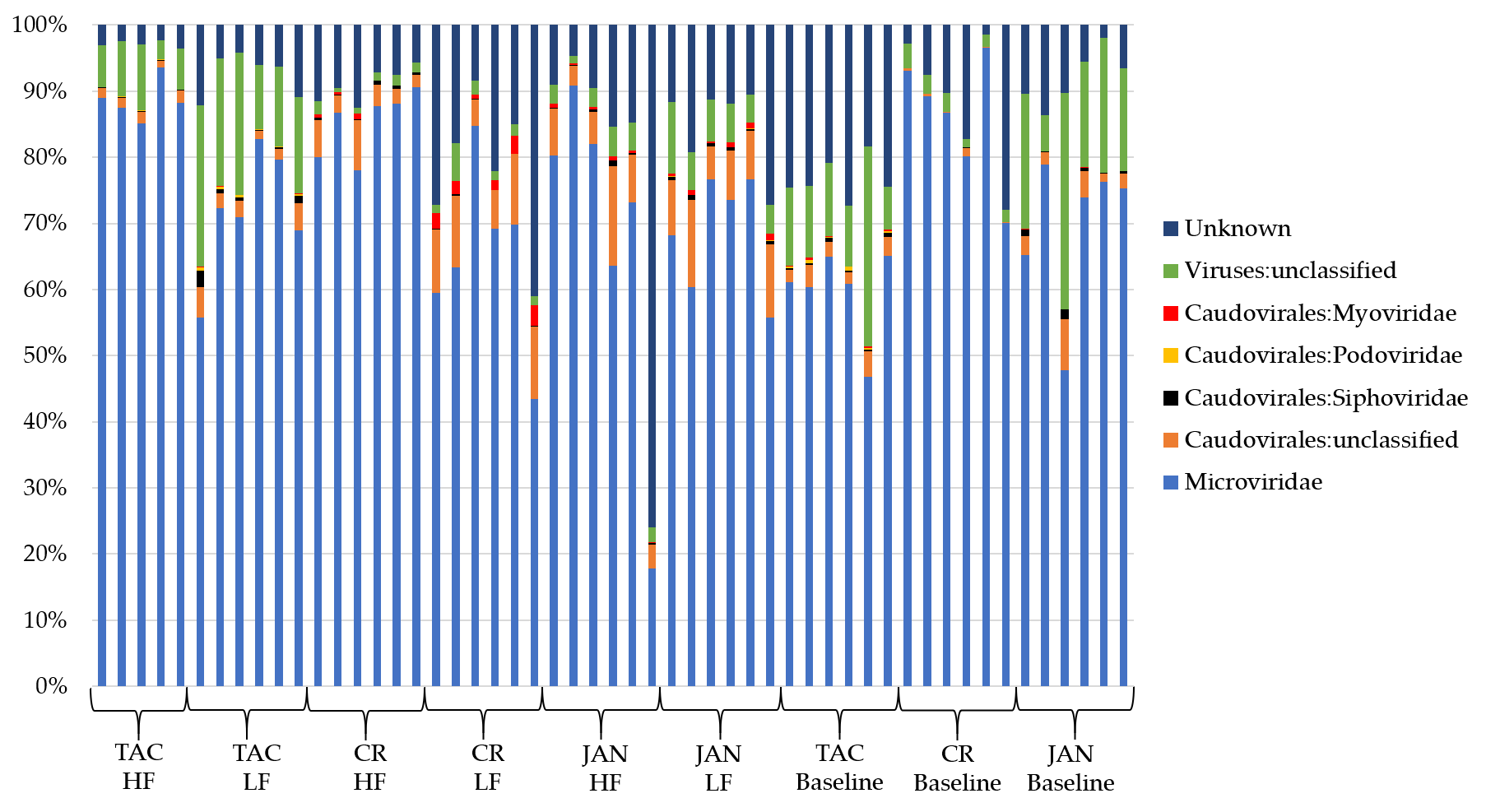

**Figure S5.** Viral taxonomy on family level of all individual cecum samples. A threshold of the relative abundance was set to 0.25%. HF = high-fat diet, LF = low-fat diet. TAC = Taconic, CR = Charles River, JAN = Janvier.

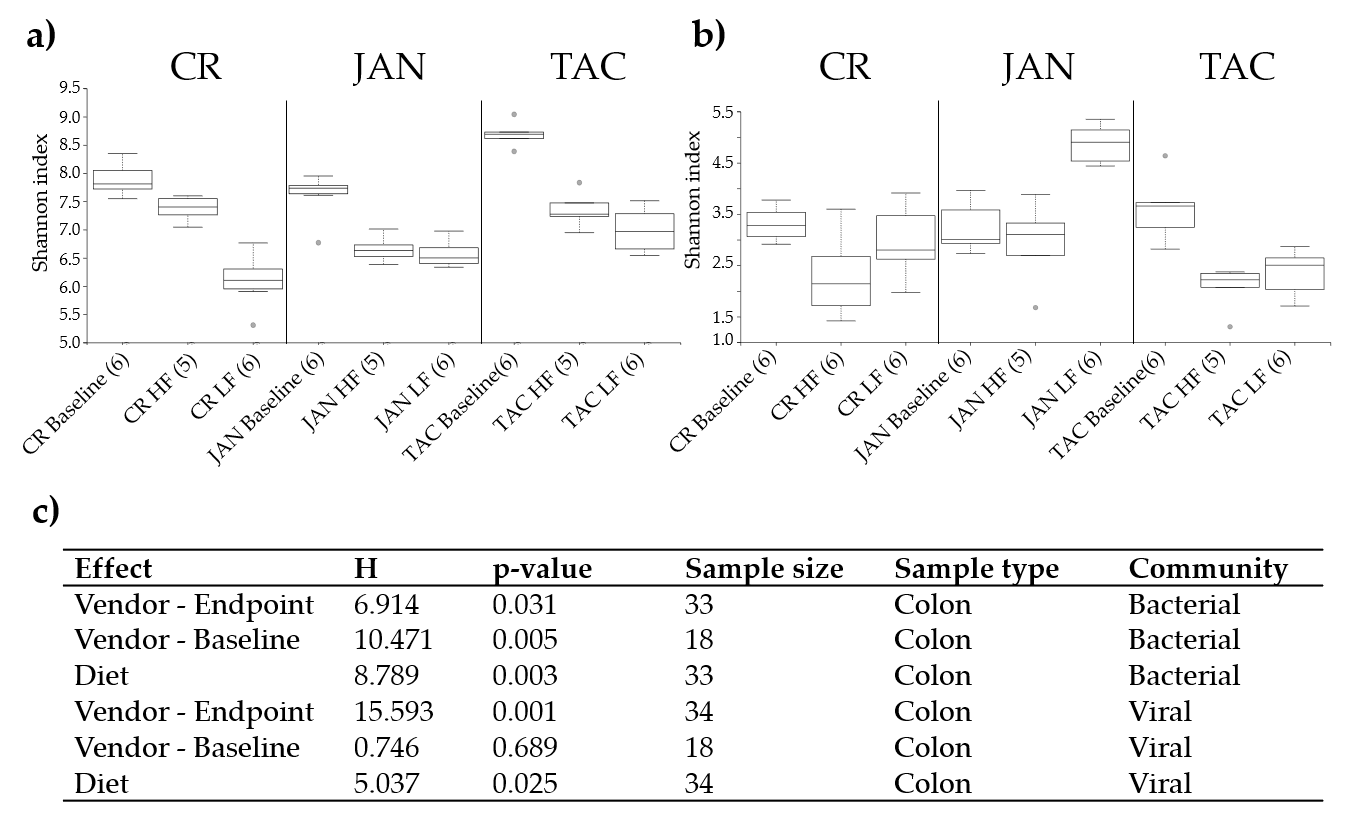

**Figure S6.** Shannon index of the colon a) bacterial and b) viral community at baseline (5 weeks of age) and after 13 weeks on low fat or high diet (18 weeks of age), respectively. The parentheses show the number of samples from each group included in the plot. c) Kruskal Wallis group analysis of the Shannon indices of the effects of diet and vendor at baseline and endpoint (18 weeks of age). Abbreviations: LF = low-fat diet, HF = high-fat diet, CR = Charles River, JAN = Janvier, TAC = Taconic.

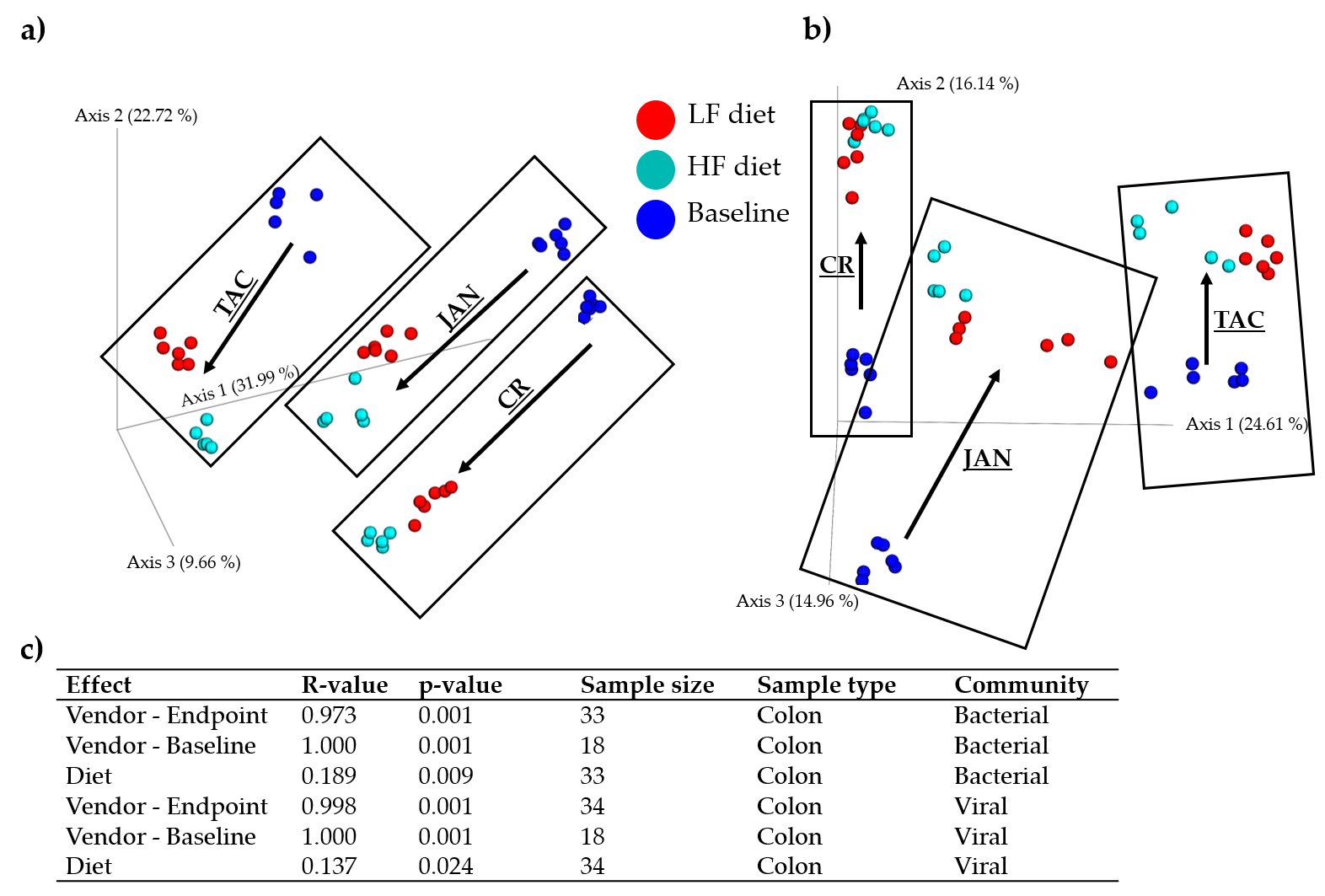

**Figure S7.** Bray Curtis dissimilarity metric PCoA based plots of a) the colon bacterial community and b) viral community at baseline (5 weeks of age) and after 13 weeks on low fat or high diet (18 weeks of age), respectively. c) ANOSIM of the Bray Curtis distances of the effects of diet and vendor at baseline and endpoint (18 weeks of age). CR = Charles River, JAN = Janvier, TAC = Taconic. Black boxes frame the samples associated to the mice vendor.

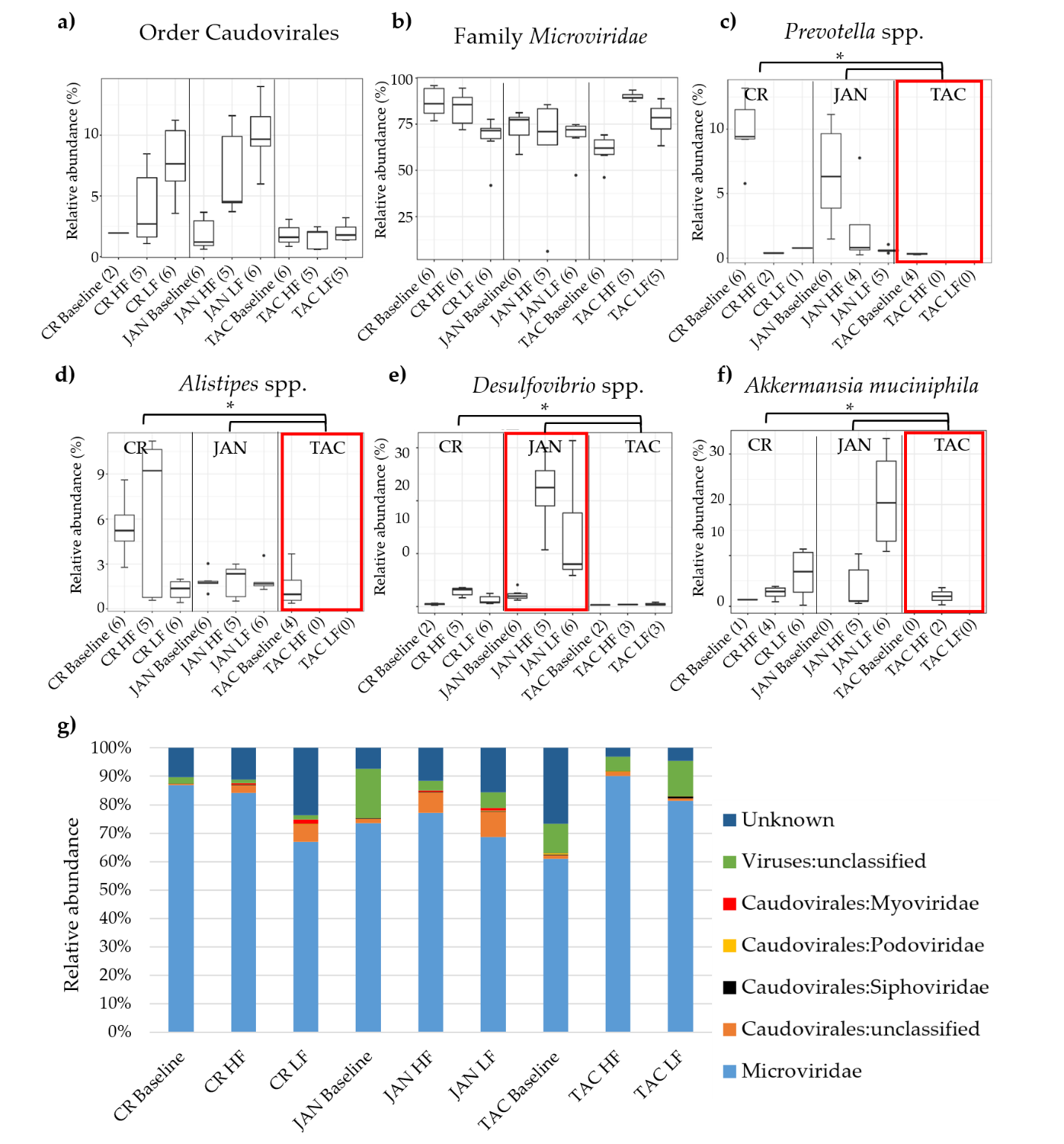

**Figure S8.** Relative abundance of a) the order Caudovirales and b) the family *Microviridae*. Differences in the relative abundance of *Prevotella* spp., *Alistipes* spp., *Desulfovibrio* spp., and *Akkermansia muciniphila* between vendors and diet are illustrated in c), d), e), and f). The most abundant viral taxonomies illustrated by bar plots in g). Black dots indicate outliers and the red boxes mark the vendor with interesting differences in bacterial abundance. Black branches and stars mark the significant bacterial differences in abundance between vendors, * = p < 0.05, ** p < 0.005, *** p < 0.0005 based on pairwise Wilcoxon rank sum test. Abbreviations: LF = low-fat diet, HF = high-fat diet, CR = Charles River, JAN = Janvier, TAC = Taconic.

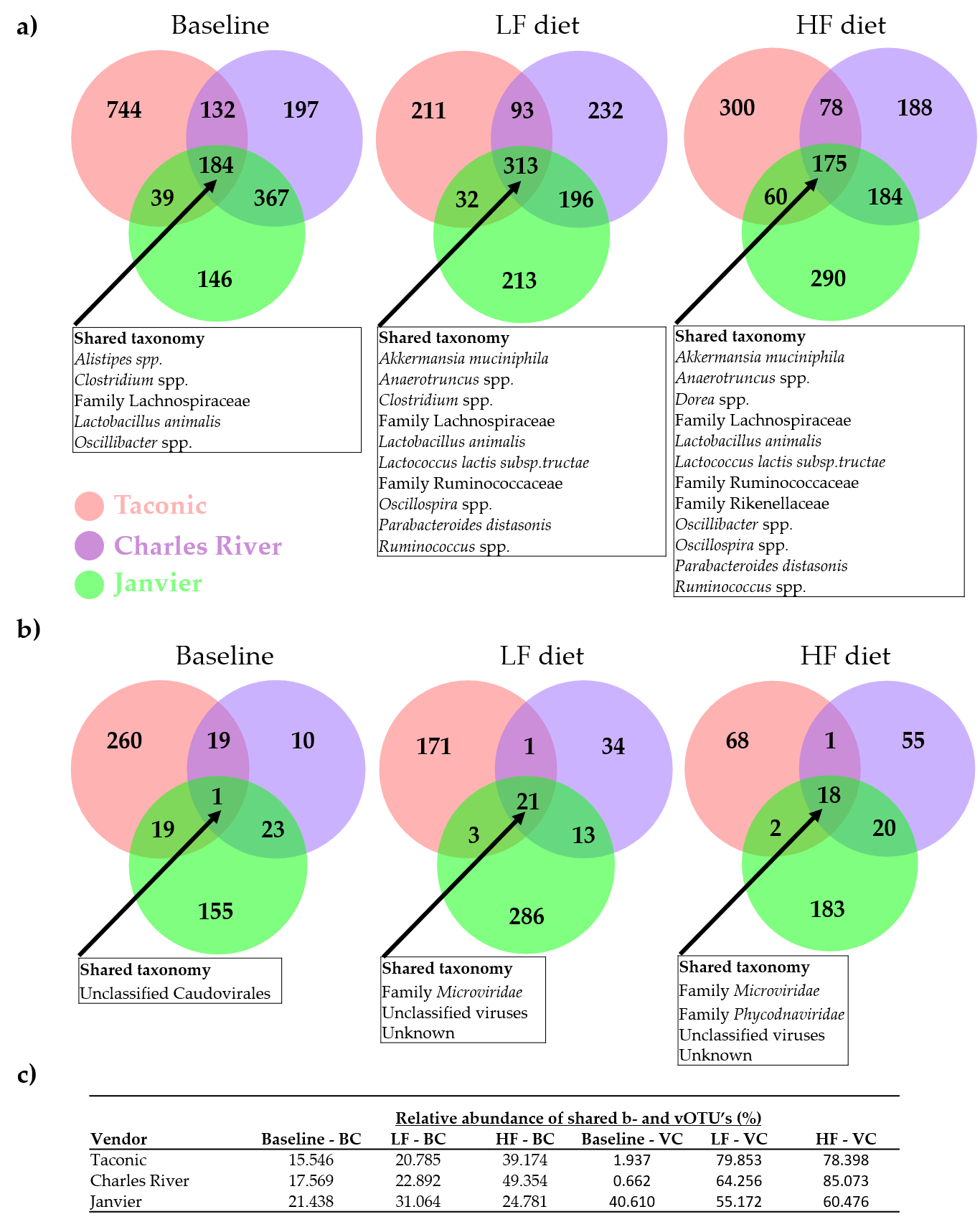

**Figure S9.** Venn diagrams illustrating the number of shared a) bacterial and b) viral OTUs (b- and vOTU’s) amongst mice purchased from three vendors at baseline (5 weeks old) and endpoint (18 weeks old) on either high-fat (HF) or low-fat (LF) diet. The boxes sum up the shared taxonomy amongst the OTU’s. c) Table showing the sum of the relative abundance of shared b- and vOTU’s from baseline to endpoint. BC = Bacterial community, VC = viral community.

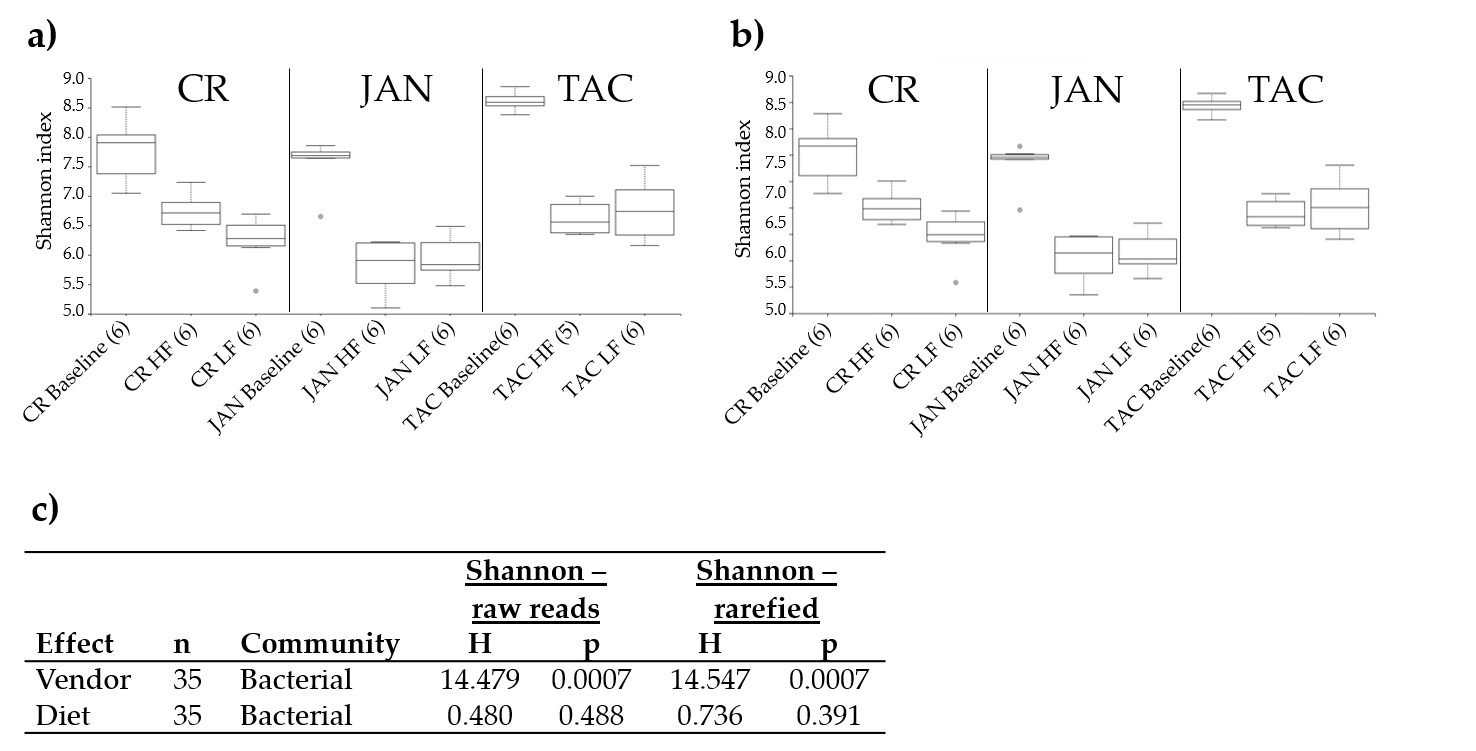

**Figure S10.** Box and whiskers plot comparing the bacterial alpha-diversity (Shannon index) when the analysis was based on either raw read count a) or rarefied read counts b). (c) Kruskal Wallis analysis emphasising that rarefaction was not necessary in this case.

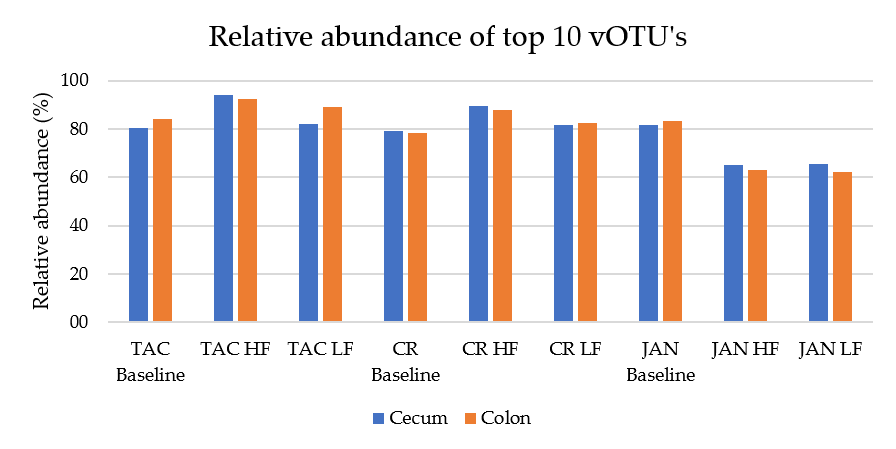

**Figure S11.** Bar plot of the mean relative abundance (%) represented by the top 10 vOTUs covering all groups of diet and vendor of both cecum and colon samples.
