## Supplementary figures and images for "Mouse vendor influence on the bacterial and viral gut composition exceeds the effect of diet"

### Figure S1.tif

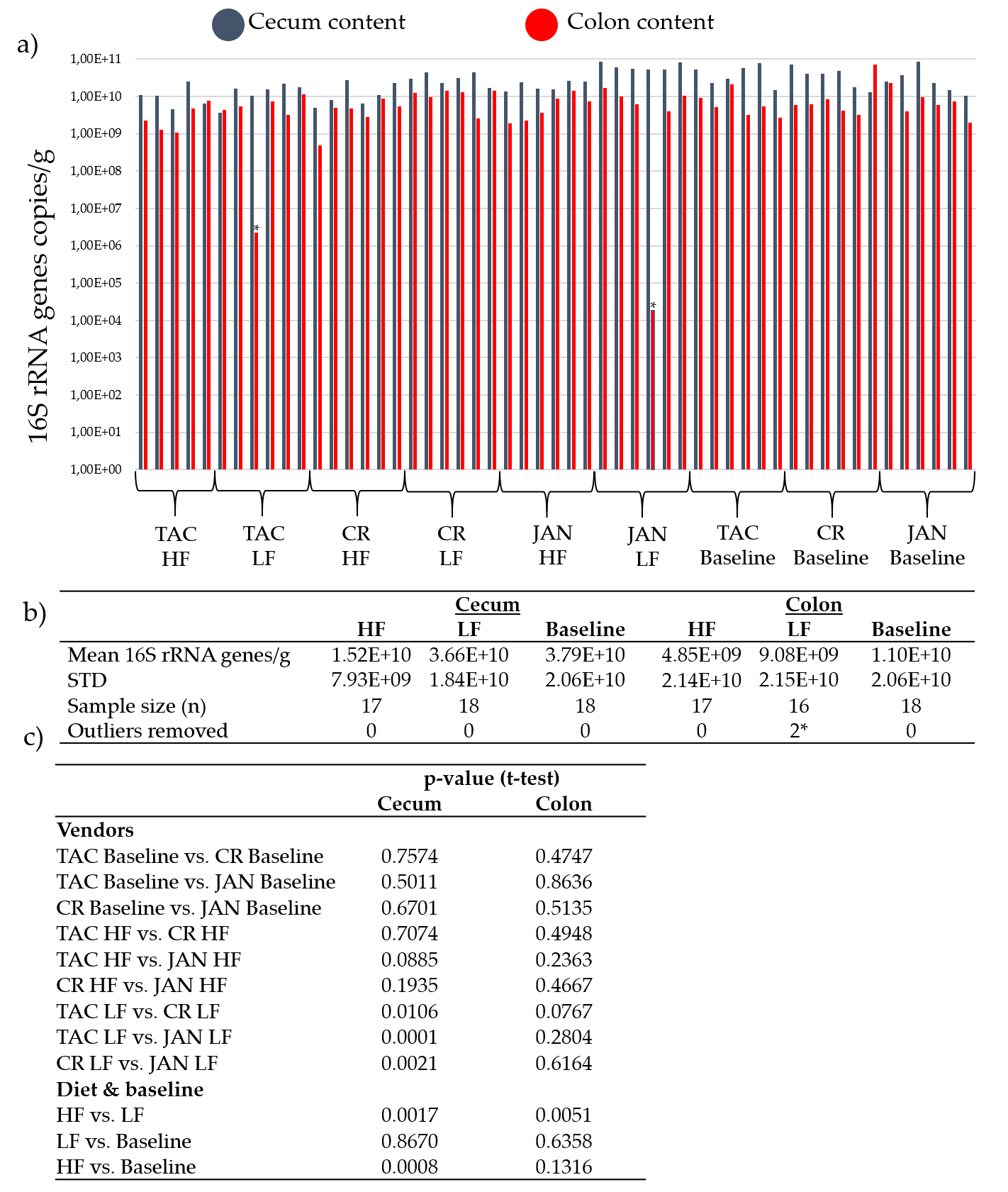

### Figure S2.tif

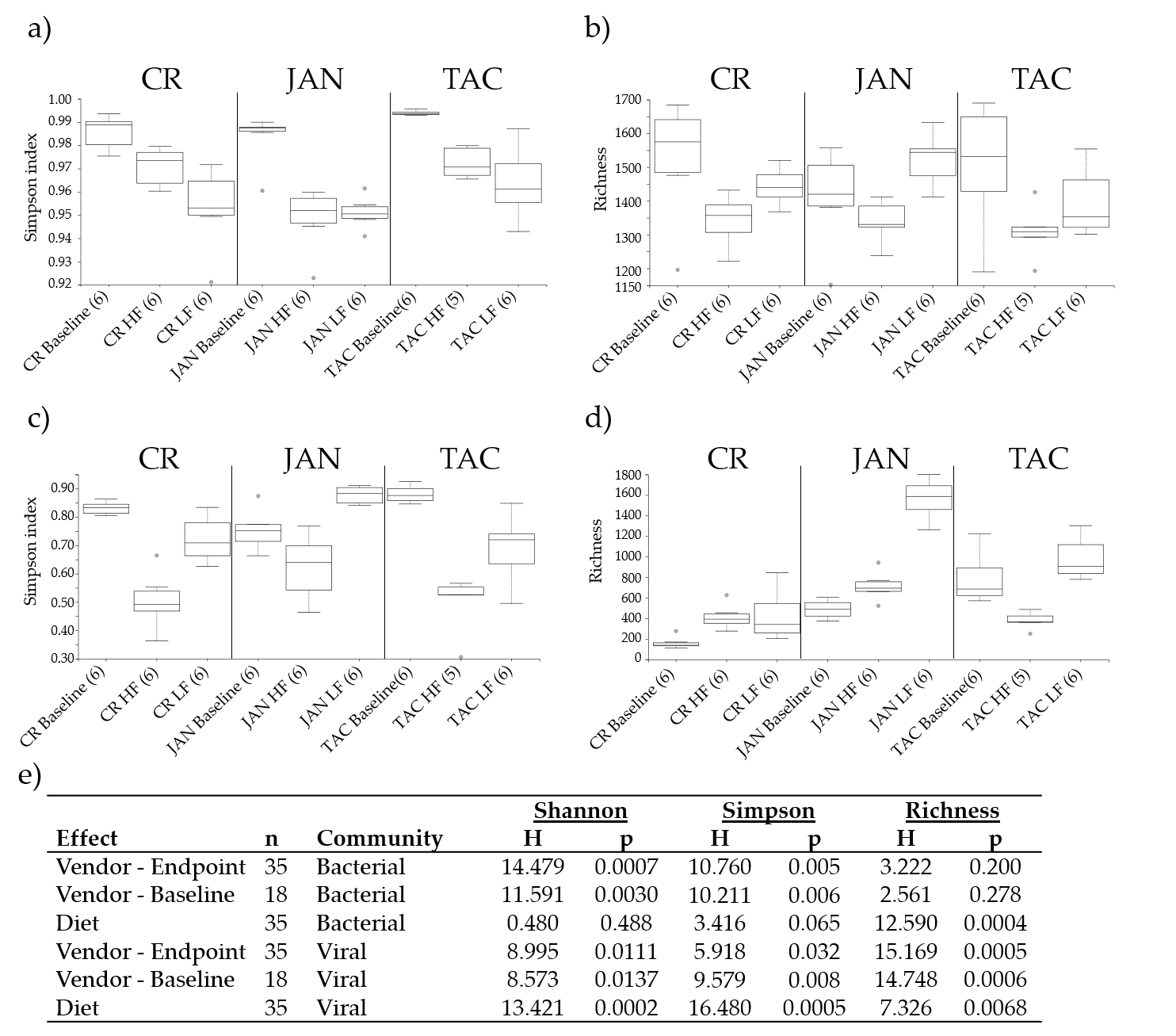

### Figure S3.tif

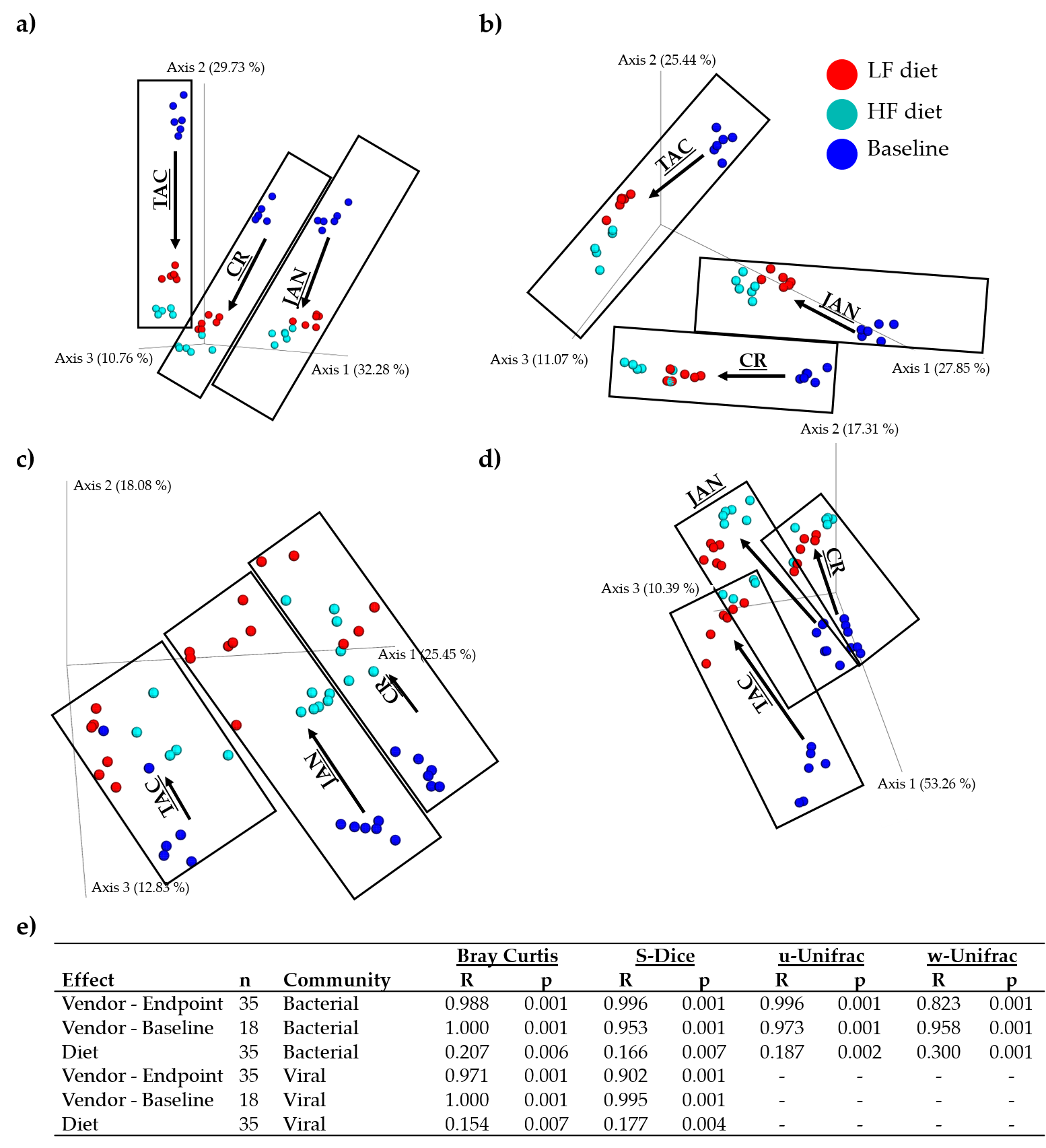

### Figure S4.tif

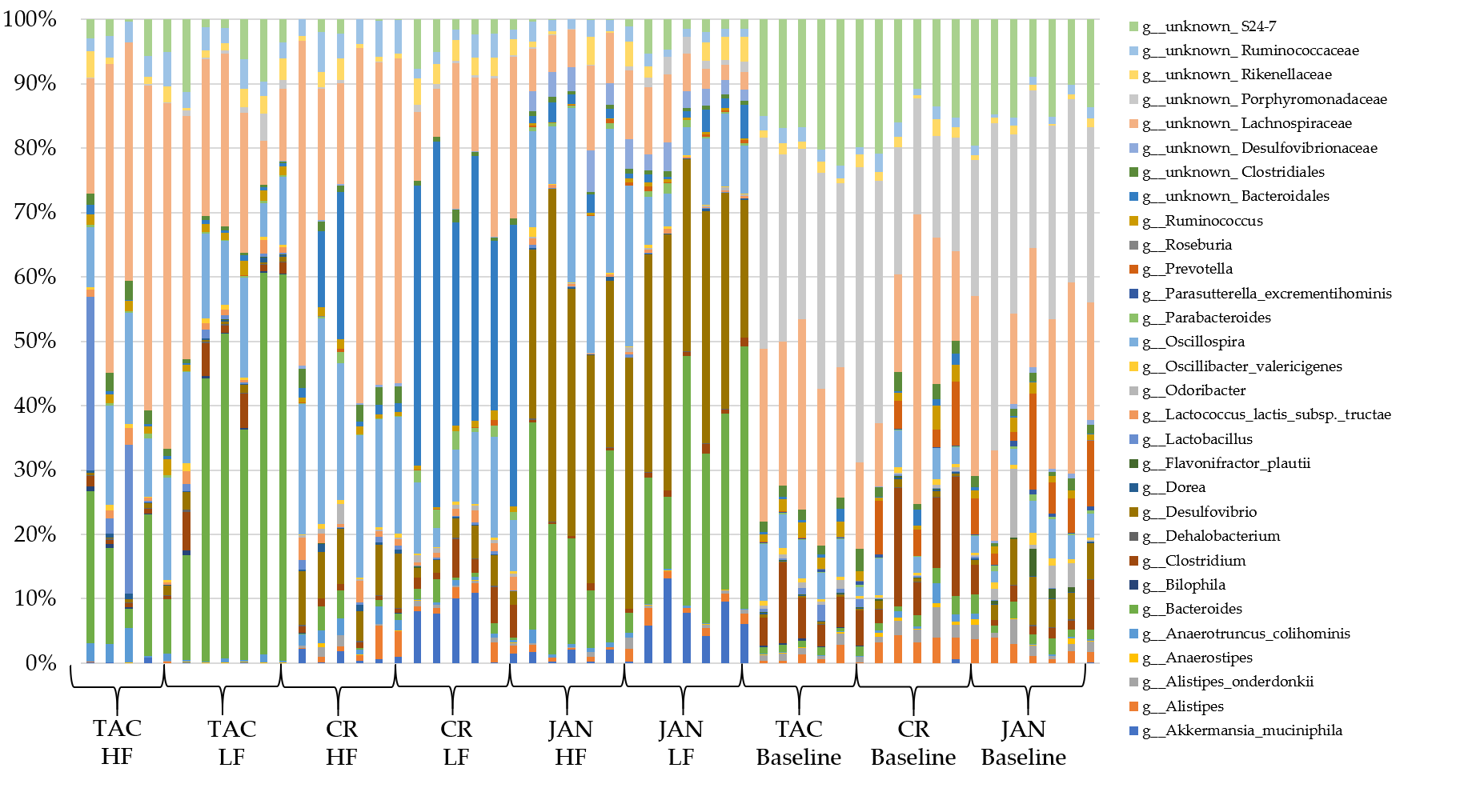

### Figure S5.tif

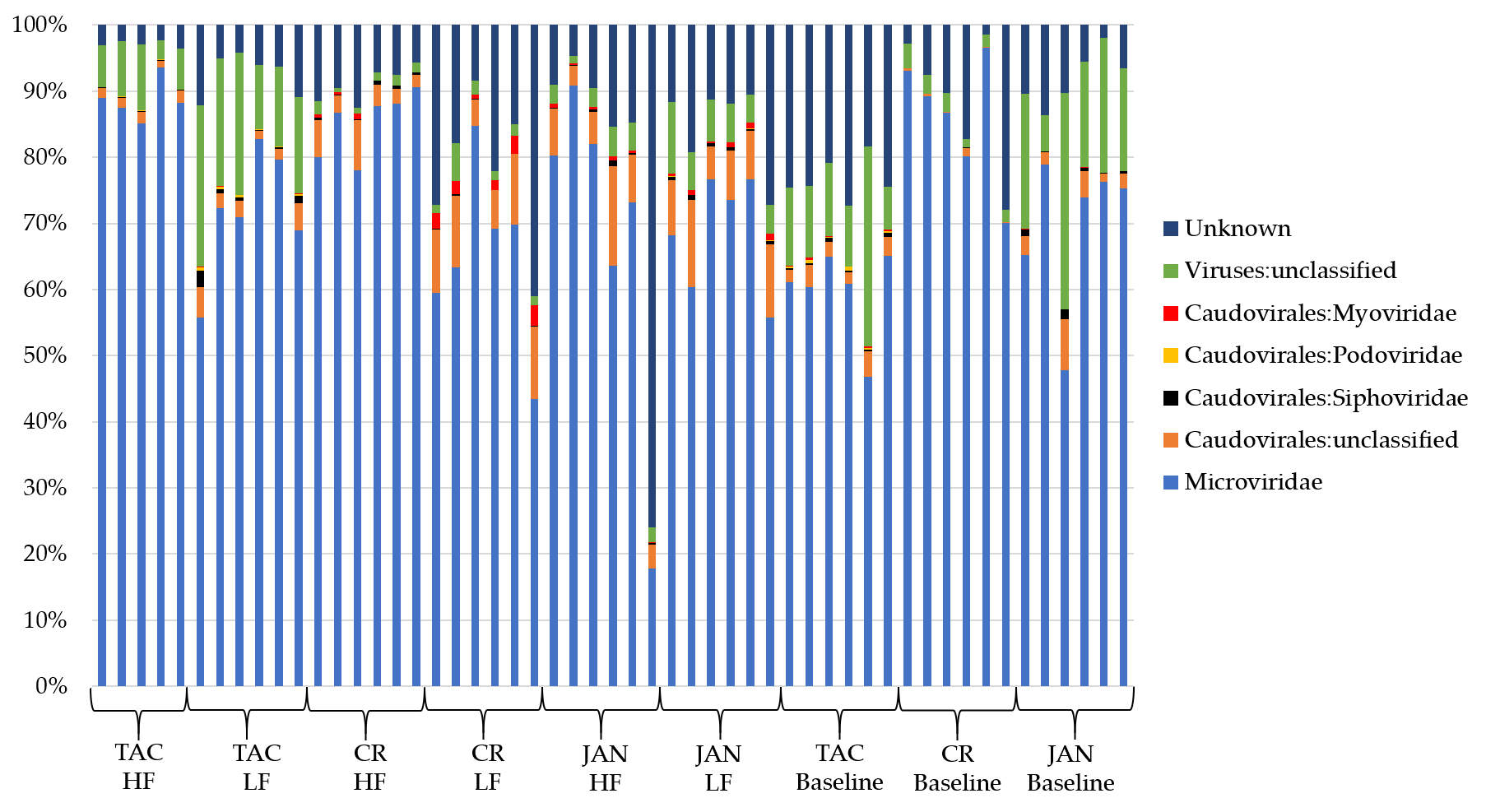

### Figure S6.tif

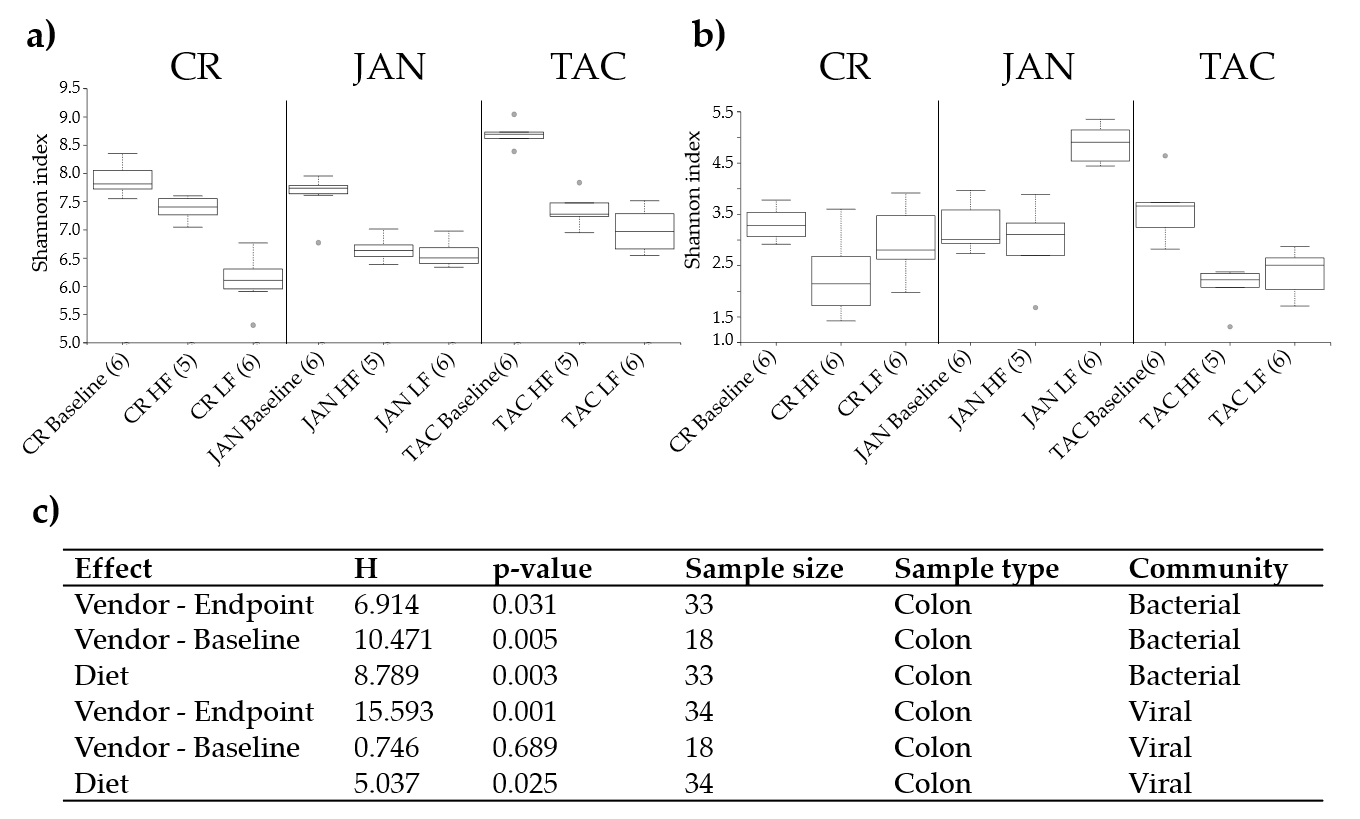

### Figure S7.tif

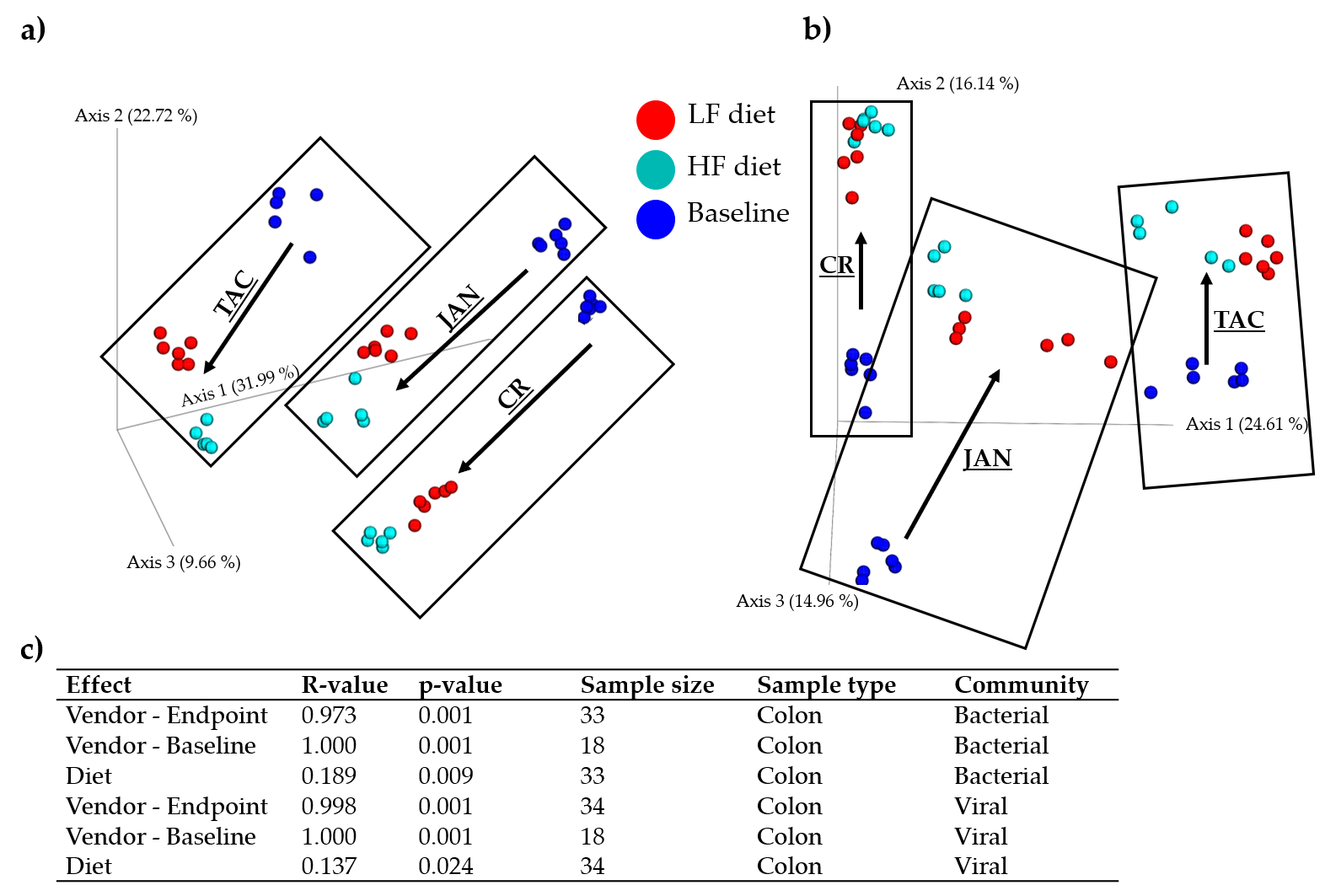

### Figure S8.tif

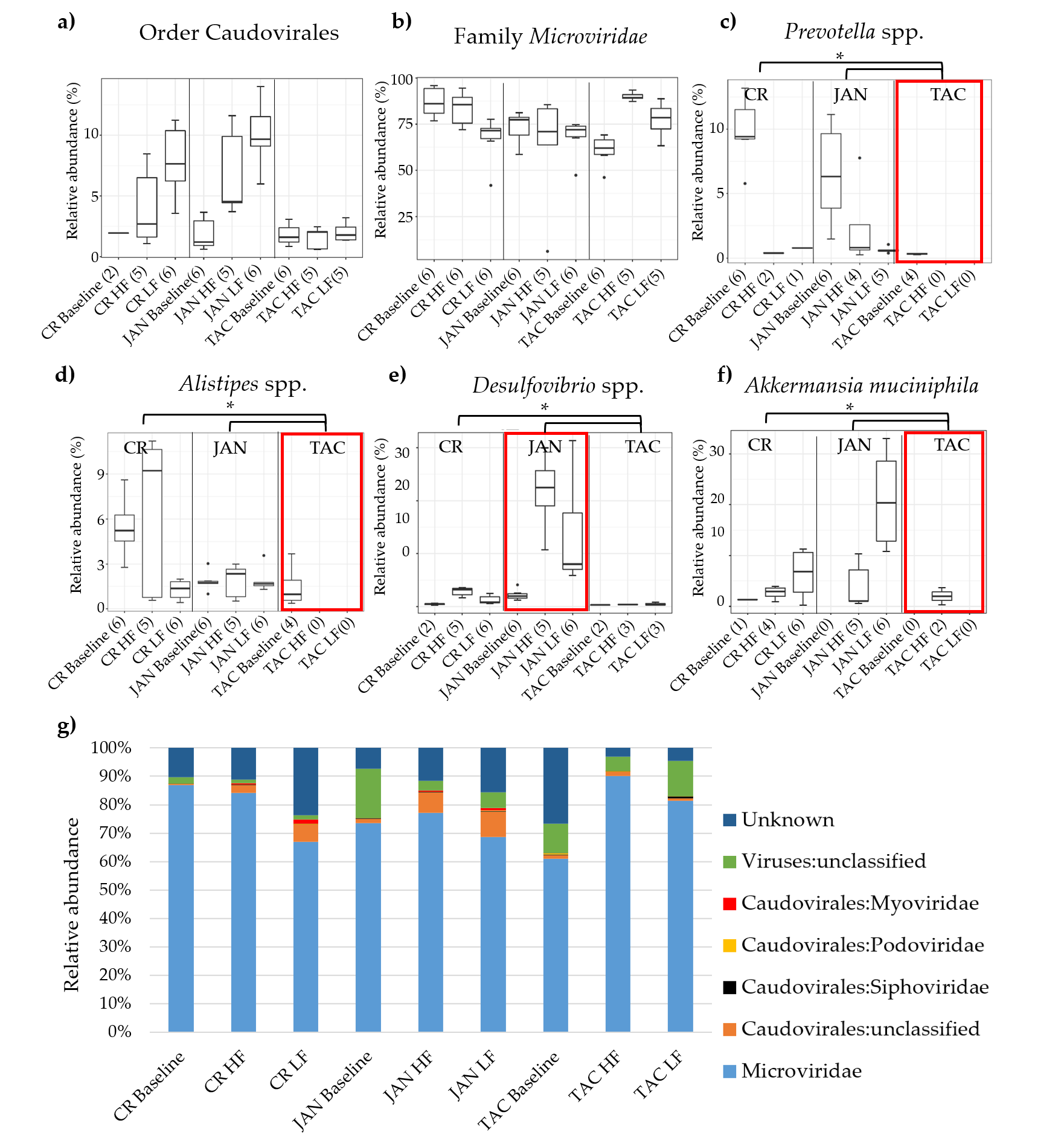

### Figure S9.tif

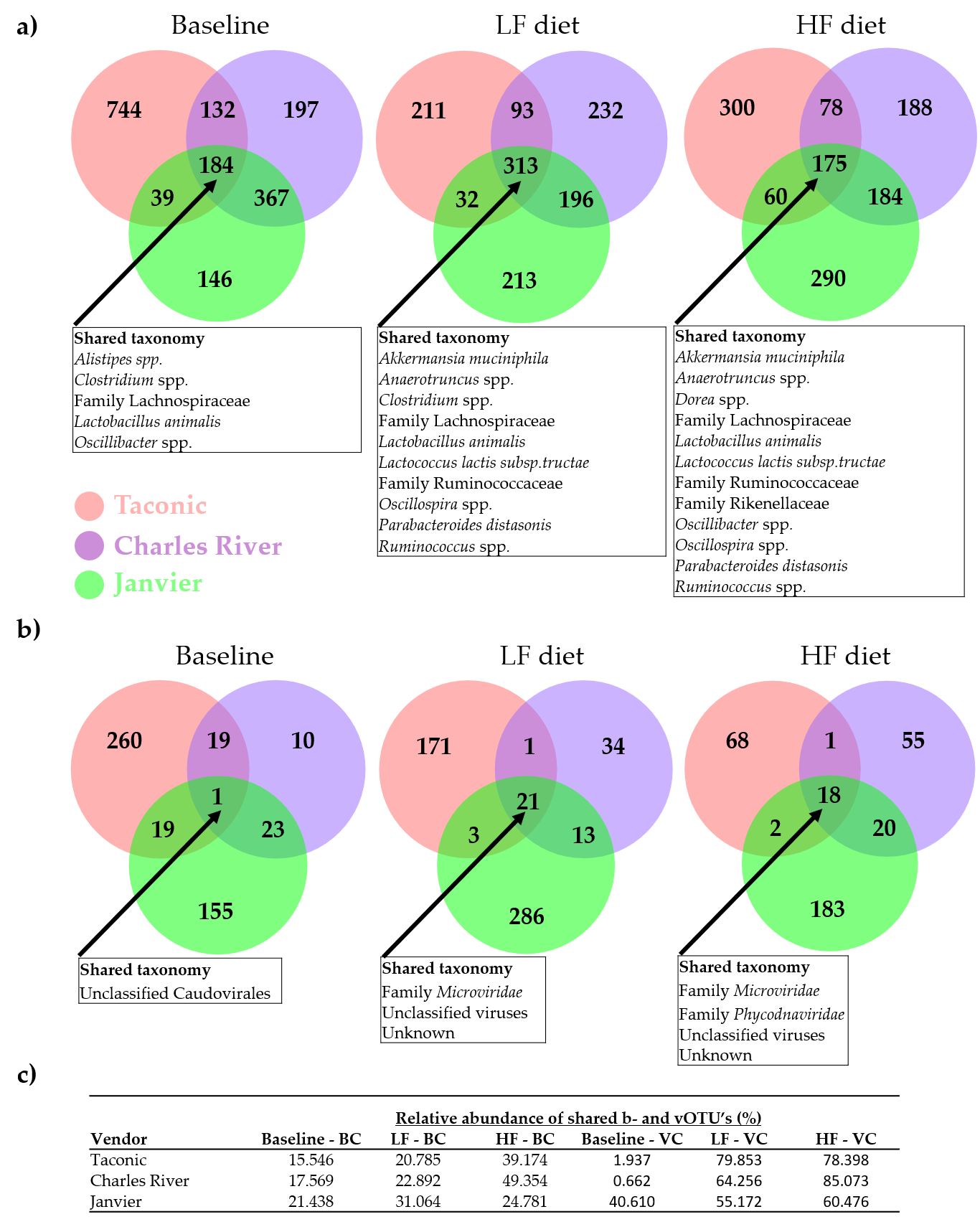

### Figure S10.tif

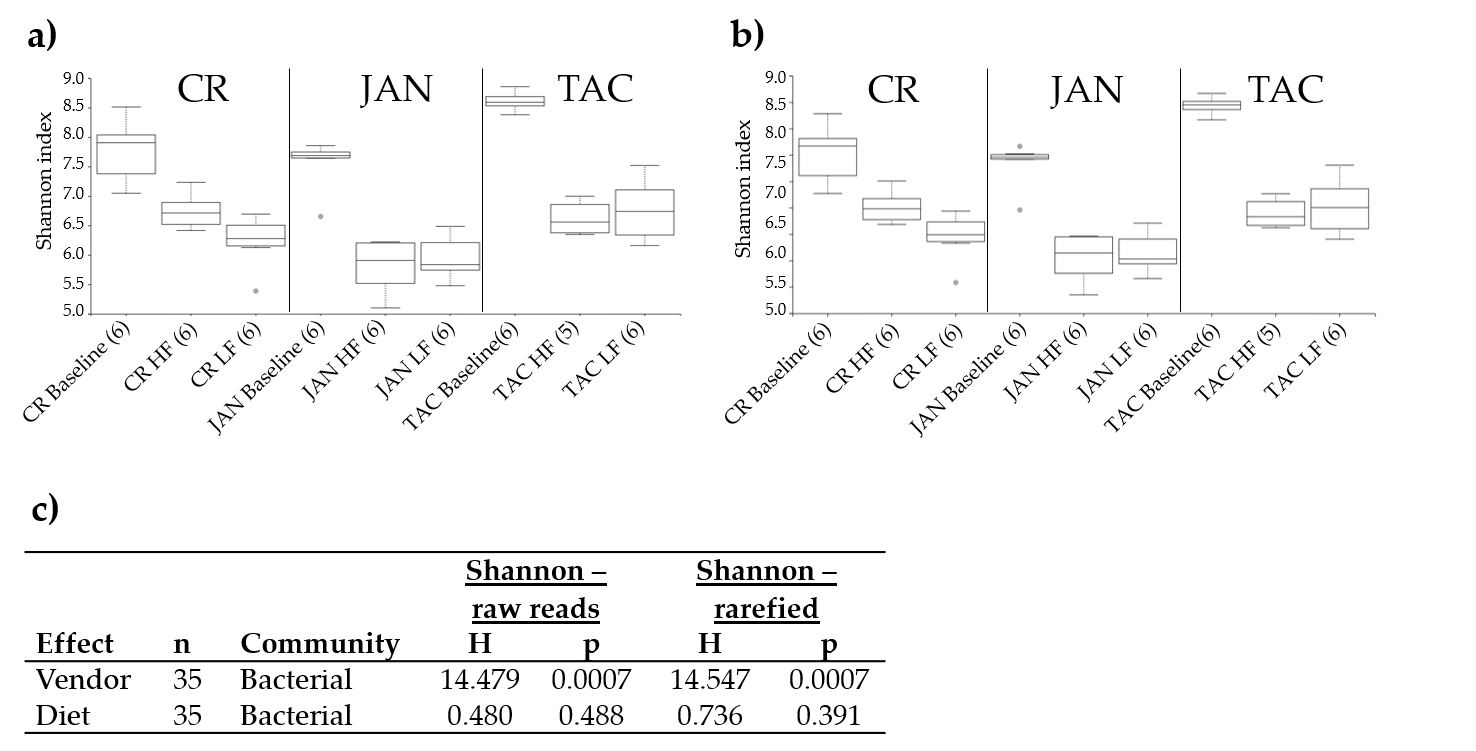

### Figure S11.tif

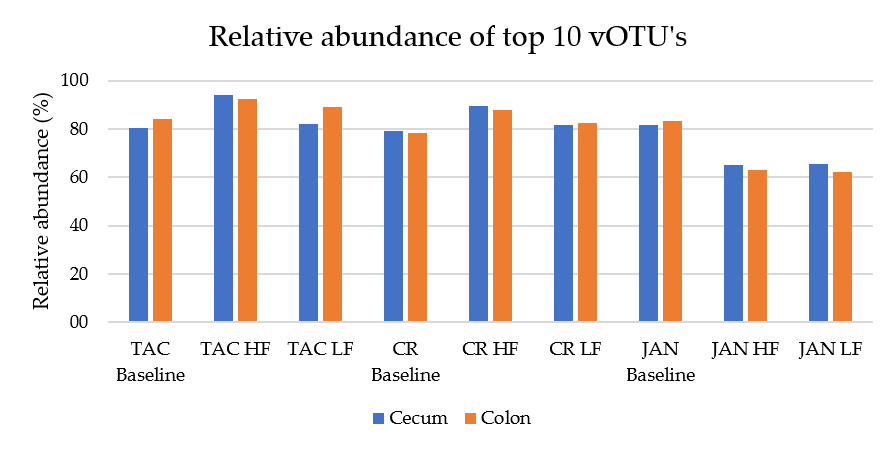
